# Rich structure landscapes in both natural and artificial RNAs revealed by mutate-and-map analysis

**DOI:** 10.1101/017624

**Authors:** Pablo Cordero, Rhiju Das

## Abstract

Landscapes exhibiting multiple secondary structures arise in natural RNA molecules that modulate gene expression, protein synthesis, and viral infection, but it is unclear whether such rich landscapes are special features of biological sequences or whether they can arise without explicit selection. We report herein that high-throughput chemical experiments can isolate an RNA’s multiple alternative secondary structures as they are stabilized by systematic mutagenesis (mutate-and-map, M^2^) and that a computational algorithm, REEFFIT, enables unbiased reconstruction of these states’ structures and populations. In an in silico benchmark on noncoding RNAs with complex landscapes, M^2^-REEFFIT recovers 95% of RNA helices present with at least 25% population while maintaining a low false discovery rate (10%) and conservative error estimates. In experimental benchmarks, M^2^-REEFFIT recovers the structure landscapes of a 35-nt MedLoop hairpin, a 110-nt 16S rRNA fourway junction with an excited state, a 25-nt bistable hairpin, and a 112-nt three-state adenine riboswitch with its expression platform, molecules whose characterization previously required expert mutational analysis and specialized NMR or chemical mapping experiments. With this validation, M^2^-REEFFIT enabled tests of whether artificial RNA sequences might exhibit complex landscapes in the absence of explicit design. An artificial flavin mononucleotide riboswitch and a randomly generated RNA sequence are found to interconvert between three or more states, including structures for which there was no design, but that could be stabilized through mutations. These results highlight the likely pervasiveness of rich landscapes with multiple secondary structures in both natural and artificial RNAs and demonstrate an automated chemical/computational route for their empirical characterization.

## INTRODUCTION

RNAs are deeply involved in gene expression, gene regulation, and structural scaffolding and are forming the basis of novel approaches to control these processes.^1-3^ Several of RNA’s natural and engineered roles rely on its ability to fold into and interconvert between multiple functional structures. Ribozymes, riboswitches, and protein-complexed RNAs transition between several states to detect and respond to small molecules and other macromolecules; to proceed through numerous steps of RNA splicing reactions; to initiate, catalyze, and proofread protein translation; to activate logical circuits in cells; and to package, release, and replicate RNA viruses.^4-9^ Rationally dissecting and re-engineering these ‘dynamic structure landscapes’ depends on knowledge of the alternative states of an RNA’s structural ensemble.^10,11^

Empirical portraits of such landscapes are missing for the vast majority of natural and engineered RNAs and it is unclear whether a rich multi-state landscape is a property specially selected by evolution or an intrinsic feature of RNA that can arise without explicit design or selection. Watson-Crick RNA secondary structure landscapes have been computationally predicted using dynamic programming techniques for decades^12,13^ and many RNAs are predicted to form multiple structures at equilibrium, with ‘non-native’ helices reaching populations of 25% or greater.^12-14^ However, difficulties in treating non-canonical interactions render these predictions inaccurate. Indeed, some studies have suggested that conformational switches are special hallmarks of biological function rather than an intrinsic feature of generic RNA sequences.^15-18^ Unfortunately, the few experimental techniques that can validate or refute multi-state models are costly and difficult. For example, single molecule methods have been successful at revealing rarely populated RNA states but have not provided enough information to infer their structures.^19^ Powerful insights have come from advances in nuclear magnetic resonance (NMR) spectroscopy^20^ but require focused technical expertise, expensive infrastructure, and RNA targets with limited structural heterogeneity. In contrast, RNA chemical mapping, or footprinting, is a simple class of techniques that can achieve single-nucleotide resolution structural data for any RNA.^21,22^ For RNAs with multiple states, however, these chemical mapping data give ensemble averages over all states, leading to a dramatic loss of information compared to what would be needed to resolve the RNA’s dynamic structure landscape and to make testable predictions.^17,23^

While developing experiments that couple systematic mutagenesis with chemical mapping (mutate-and-map, or M^2^), we have observed that single mutations can produce dramatic changes in chemical mapping data throughout an RNA sequence, with several mutations often giving the same alternative pattern.^24,25^ We hypothesized that these perturbed patterns correspond to the reweighting of the structural ensemble of the RNA so that alternative component states dominate the chemical mapping data. We have recently shown that such alternative structures can be inferred after expert inspection of M^2^ measurements and extensive compensatory rescue studies in an *E. coli* 16S ribosomal RNA domain.^26^ Reasoning that such landscape dissection might be fully automated through the use of blind source separation algorithms, we have now developed an analysis framework called the RNA Ensemble Extraction From Footprinting Insights Technique (REEFFIT; see Supporting Figure S1).

Here, we sought to use M^2^-REEFFIT to determine whether complex landscapes could arise in artificial RNAs without explicit natural selection or design. Prior techniques for detecting alternative structures, including covariation-based methods^16^ and the recent RING-MaP method ^17^, have not been benchmarked in cases with well-characterized landscapes and may be biased towards or against alternative structures. Therefore, we developed three tests for M^2^-REEFFIT landscape dissection. First, a benchmark on simulated M^2^ data for 20 natural non-coding RNA sequences provided ‘gold standard’ reference results allowing unambiguous assessment of accuracy. Second, we applied M^2^-REEFFIT to four biological and artificial RNAs that had been previously characterized in detail by NMR or chemical mapping and recovered the landscapes defined through prior expert analysis. Third, we developed a validation approach based on stabilizing and experimentally testing predicted structures by multiple mutations. With these benchmarks and methodological developments, we used M^2^- REEFFIT to demonstrate that an imperfectly designed riboswitch and a randomly generated RNA sequence each form at least three structures, including states that could not be predicted from computational modeling alone.

## RESULTS

### Dissecting landscapes de novo with M^2^-REEFFIT

We developed REEFFIT as a tool for simultaneously inferring multiple structures and their population fractions across the mutant RNAs in M^2^ measurements (see Supporting Methods for formal description, optimization schemes tested, bootstrapping, and cross-validation). To test its accuracy in a setting with unambiguous ground truth, we first applied M^2^-REEFFIT to infer structure landscapes for an *in silico* benchmark of 20 sequences drawn from the RFAM database of non-coding RNA molecules. We simulated M^2^ data by randomly choosing up to 200 suboptimal structures in the ensembles of the wild type sequence and of all variants that mutate a single nucleotide to its complement, as are typically probed in M^2^ experiments. We then simulated these structures’ respective SHAPE reactivity profiles using known reactivity distributions.^27^ To mimic inaccuracies in available energetic models, we re-weighted the ensemble randomly by introducing Gaussian noise to the predicted free energies. Our simulated benchmark for landscape modeling included RNAs that folded into predominantly one structure, RNAs with excited states present at more than 10% population, and RNAs with complex landscapes in which various competing helices were present. Below we describe examples of each of these cases.

Figures 1a-e show simulated data for the sequence RF01301, the small nucleolar RNA SNOR4A. The benchmark simulation of this RNA involved 25 suboptimal secondary structures, most of them related by register-shifts to either a 10 nucleotide hairpin, 1301-A, or a more complex structure 1301-B comprised of 5 helices separated by internal loops (Figure 1e). The REEFFIT model provided an excellent fit to this RNA’s simulated M^2^ data (Figures 1a and 1b). Recovered features included the ‘diagonal’ feature corresponding to disruptions near the site of each mutation as well as more dramatic features due to stabilization of state 1301-B upon certain mutations, visible as ‘plaid’ patterns in the simulated data (see, e.g. U14A, A41U). The importance of using a full ensemble fit was tested by repeating REEFFIT with the constraint that only 1, 2, or 3 structures were used in describing the data; these fits did not adequately capture all features in the simulated M^2^ data (Supporting Figure S3).

**Figure 1:**
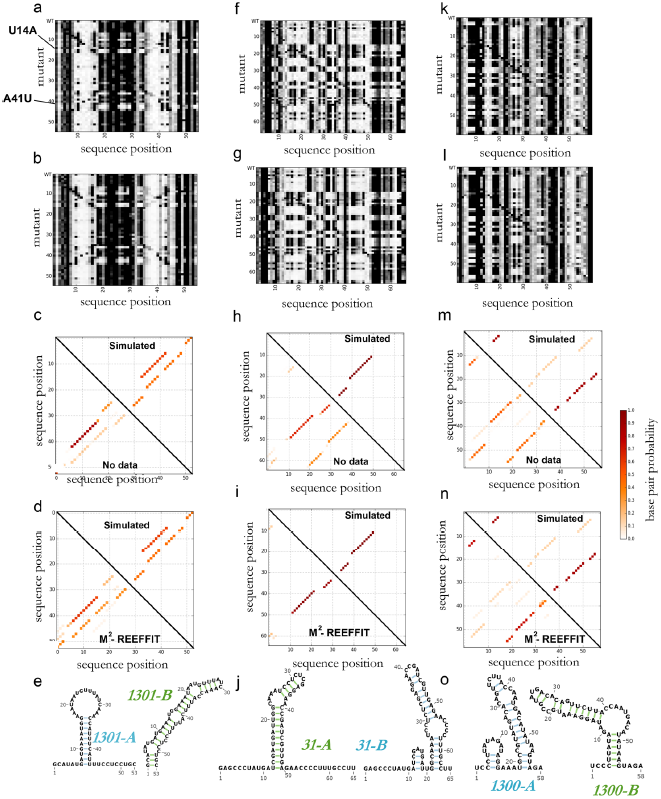
Three example cases of evaluating REEFFIT with simulated data. (a) Simulated M^2^ data for the RF01301 snoR4a test case (b) Bootstrapped REEFFIt fits. (c) Base pair probability matrix comparisons between the simulated (true) base pair probabilities (upper half) and the unperturbed RNAstructure values; (d) analogous base pair probability comparison to M^2^-REEFFIT estimates. (e) Cluster medioids of structures used for generating the M^2^ simulated data. (f—j) Evaluation results for the RF00031 SECIS RNA simulated mapping data. (k—o) Evaluation results for the RF00014 DrsA RNA.

Beyond the quality of the fit, the actual details of the ensemble were recovered accurately, despite a substantial perturbation of states’ population fractions from their original RNAstructure values. The improvement can be visualized in the matrices of base pair probabilities from RNAstructure and from REEFFIT compared to the actual simulated matrix (Figure 1c and 1d). Quantitatively, the total weights for 1301-A and 1301-B in the wild type sequence, simulated to be 58% and 42%, respectively, were recovered as helix fractions of 62±12% and 38±11%, respectively. Analogous accuracy was observed across all the simulated mutants (see Supporting Data). Having the full mutate-and-map data set was important for making these inferences; using the wild type data alone (as would be carried out in conventional ‘one-dimensional’ chemical mapping ^23^) returns a highly ambiguous fit with incorrect helix frequencies (see Supporting Figure S4).

Figures 1f-o illustrate REEFFIT analyses for two other benchmark RNAs with qualitatively distinct ensembles. Tests on simulated data for the selenocysteine insertion sequence (SECIS, Rfam id RF00031, Figs. 1f-j) supported the ability of M^2^-REEFFIT to model an RNA with a single dominant secondary structure (95% population, recovered as 91±15%) without giving false positives for alternative structures present at low population. For complex cases, such as RF01300, REEFFIT was able to detect the 190 helices present at more than 25% population in the multiple states of the simulated ensembles of the wild type and 58 mutants (Fig. 1k-o). In both of these examples, analysis of the data for the wild type sequence alone gave poor recovery of the simulated ensembles (see Supporting Figure S4), confirming the need for the full M^2^ measurements.

Over the entire 20-RNA benchmark, REEFFIT was able to consistently detect the presence of dominant and alternative helices (Supporting Results and Supporting Figures S2-S4). We set a criterion for helix detection that the fitted population should be larger than the population error estimated from bootstrapping, i.e. the signal-to-noise ratio should be greater than one. With this criterion, REEFFIT achieved 94.6% sensitivity over helices present with at least 3 base pairs and at least 25% population, corresponding to a false negative rate (FNR) of 5.4%. The false discovery rate (FDR) was low as well, at 9.7% (see Table 1 and Supporting Table S1; full precision-recall curves given in Supporting Figure S2). Without data, the error rates were substantially worse, by three-fold and two-fold, respectively (FNR of 18.0% and FDR of 23.3%). Values for base-pair-level error rates were similar to helix-level error rates (Supporting Table S2). Error rates for wild type sequences alone (i.e., excluding singlenucleotide mutants) were higher but also showed a strong improvement in REEFFIT ensembles compared to ensembles modeled with no data (FNR of 15.2%, FDR of 15.2% for REEFFIT; FNR of 24.5% and FDR of 25.4% without data, see Supporting Table S3). Use of only wild type data and no mutants, as would be carried out in conventional chemical mapping measurements, gave high error rates similar to landscapes modeled without data (FNR of 22.7% and FDR of 25%, see Supporting Information Table S4 and an example in Supporting Results and Supporting Figure S4), confirming the necessity of M^2^ data for accurate landscape dissection. Further systematic checks and use of ViennaRNA^28^ initial models and benchmark evaluation using helix-wise RMSD are given in the Supporting Results and Supporting Tables S5, S6, and S7.

**Table 1:**
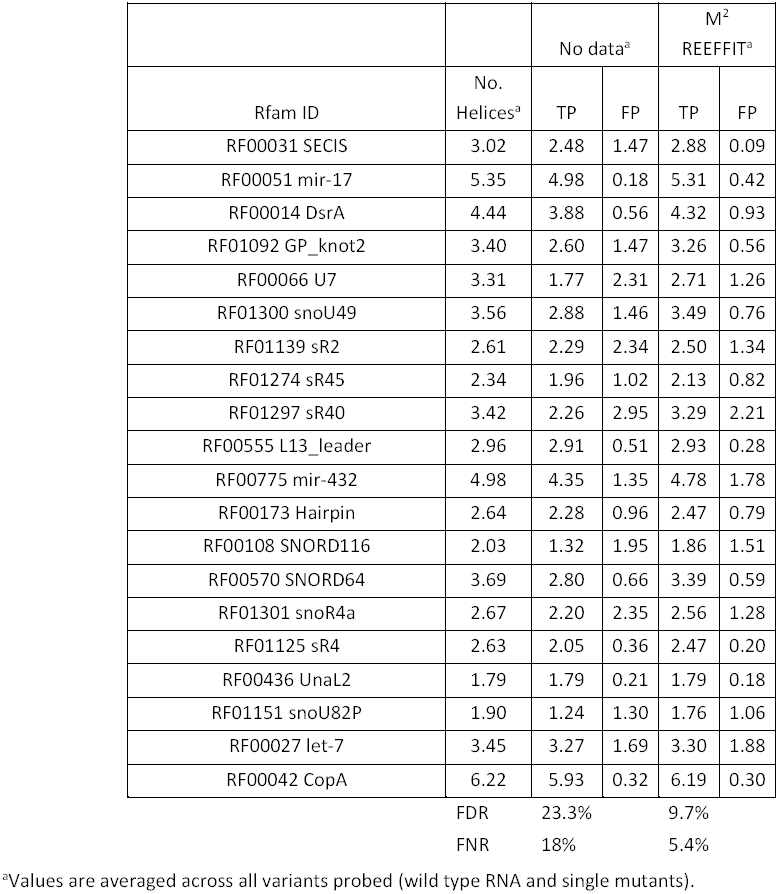
Table 1: Performance results for M^2^-REEFFIT landscape dissection on a 20 RNA benchmark from the RFAM database.

### Experimental tests on RNAs previously characterized by M^2^ and expert analysis

Success in the above *in silico* benchmark suggested that REEFFIT would accurately recover prior analyses of experimental M^2^ data sets, such as the MedLoop RNA (Figure 2a-d). This RNA was previously designed to exhibit a single, stable 10 base pair helix with a 15 nucleotide loop (MLP-A, Figure 2d).^24^ A few mutations were observed to give a clearly distinct chemical mapping pattern and predicted computationally to fold into an alternative structure (MLP-B in Figure 2a; see e.g. G4C in M^2^ data) but not expected to be strongly populated in the wild type sequence. Automated REEFFIT modeling of the MedLoop M^2^ data recovered both the dominant structure MLP-A and the alternative state MLP-B, and bootstrapped uncertainties gave bounds on the frequency of MLP-B in the wild-type sequence (4±4 %, see mutant-wise state fractions in Supporting Figures S6a-c). Thus, REEFFIT was able to explain rearrangements of mutants into an alternative state, but at the same time did not overpredict its presence in the wild-type sequence (Figure 2c).

**Figure 2:**
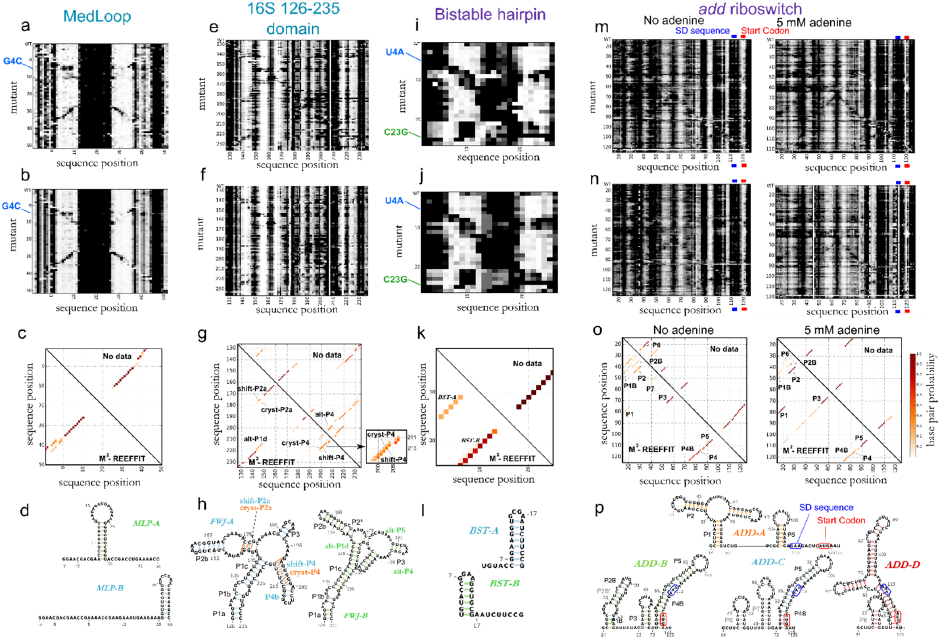
Landscape dissection of four diverse RNA systems from experimental mutate-and-map (M^2^) data. (a) M^2^ measurements for the MedLoop artificial sequence (given to REEFFIT as input); (b) REEFFIT bootstrapped fits; (c) base pair probability matrices of the structures in the wild type landscape (upper triangle: base pair probability values using no data, lower triangle: REEFFIT calculated base pair probabilities); and (d) structure medioids found *de novo* by REEFFIT. M^2^-REEFFIT analysis are also presented for (e-h) a four-way junction domain of the 16S rRNA, (i-l) an artificial bistable hairpin, (m-p) and a segment of the *V. Vulnificus add* adenine riboswitch in 0 and 5 mM adenine. In (d), The Shine-Dalgarno (SD) ribosome-binding sequence was observed to be SHAPE-reactive even in the absence of adenine; while consistent with this region’s unpaired status without and with ligand in prior NMR studies, our studies indicate a more dramatic switch in this region for more complete *add* riboswitch constructs and will be reported elsewhere. See Supporting Tables S8-9 for RNA sequences and fit summaries.

As a more difficult test, we applied the algorithm to the 126-235 region of the *E. coli* 16S ribosomal RNA. In protein-bound ribosome crystals, this domain forms a four-way junction with helices P1a-c, P2a-b, P2*, P3, and P4a-b, but its solution state has been controversial^26^ (Fig. 2e-h). Conventional SHAPE-guided analysis suggested loss of P2a and P4, and formation of alternative helices alt-P1d and alt-P ^29^ (Fig. 2h, green structure). However, M^2^ and compensatory rescue experiments gave no evidence for the SHAPE-based model but instead recovered the dominant solution structure to be the holo-ribosome conformation, except for a register shift in P4a (called shift-P4a); the crystallographic P4 register was also detected as a 20±10% ‘excited state’.^26^ In agreement with this detailed analysis, automated REEFFIT analysis of the 126-235 RNA M^2^ data also returned helices P1a-c, P2*, P2b, and P3 with high population fractions (>80%, Fig. 2g). Importantly, REEFFIT recovered an admixture of P4 (21±16%) and shift-P4 (60±23%) in a 3:1 ratio, in agreement with prior analysis (Fig 2g, magnification); RNAstructure calculations or use of wild type SHAPE data alone assigned negligible probability or highly uncertain population fractions to these helices, respectively (Fig. 2g). The population of alternative helices alt-P1d and alt-P4 were found to have low populations and high errors (see Supporting Results). Further, a refined REEFFIT analysis including data for compensatory rescue double mutants recovered, with conservative error estimates, prior expert analysis (see Supporting Results and Supporting Figure S5), illustrating the automation of modeling of even a complex RNA structure landscape.

### Experimental tests on RNAs with NMR-characterized landscapes

Encouraged by REEFFIT’s performance in previous test cases for chemical mapping, we sought tests involving fully independent experimental characterization by NMR. We first investigated a bistable RNA sequence whose landscape was dissected by Höbartner and colleagues by decomposing its NH…N ^1^H NMR spectrum into a weighted sum of two states forming different hairpins (here called BST-A and BST-B).^30^ As expected, M^2^ measurements gave clear evidence for two distinct chemical mapping profiles, reflecting the bistable nature of this RNA (Figure 2i-l). One profile, consistent with BST-A appeared in variants with mutations in the 5’ end of the sequence, such as U4A. Another profile, consistent with BST-B, was revealed by mutations in the 3’ end such as C23G, which would destabilize the BST-A hairpin (Fig 2i). Both structures were recovered by REEFFIT analysis, with population weights of 73 ± 11 % and 26 ± 9 %, for BST-A and BST-B respectively, in agreement with the previously reported fractions measured by NMR (70 ± 5% and 30 ± 5 %, respectively) and correcting the erroneous weights predicted with no data (99% and <1%, respectively).

We then challenged M^2^-REEFFIT with a more complex test case: a *Vibrio vulnificus* adenosine deaminase *(add)* mRNA riboswitch (Fig. 2m-p) that, in response to the ligand adenine, exposes start site segments (Shine Dalgarno sequence and/or AUG start codon) to promote mRNA translation. In a detailed NMR study, spectra in ligand-free conditions fit well to a model with two states, apoA (∼30%, with helices P1, P2, P3, P4, and P5) and apoB (∼70%, with P1B, P2B, P3, an extended P4 called P4B, and P5). Addition of adenine ligand resulted in spectra dominated by a state holo with P1, P2, P3, and P5 and perturbed chemical shifts consistent with adenine binding to the aptamer. ^31^ While three states gave the simplest model for the prior data, more complex multistate models were consistent as well and would be generally predicted from RNA secondary structure calculations.^12,13^ We focused on whether M^2^-REEFFIT could recover the base pairs of P1, P2, P1B, P2B, P3, P4, P4B, and P5, which were unambiguously determined through NOE spectroscopy and model construct comparisons. Even in the absence of adenine, our M^2^ measurements of the RNA suggested the presence of at least two distinct structures that protect or expose the mRNA start site (AUG at nts 120-122) and, in an anticorrelated manner, expose or protect segments in the aptamer region (e.g., nts 53-60 and 66-72), respectively (M^2^ data in Fig. 2m). In addition to observing these different states upon mutation, addition of 5 mM adenine induced clear changes in the M^2^ data, including protections in the aptamer and consistent exposure of the AUG start codon in most mutants (M^2^ data in Fig. 2m, bottom; Shine-Dalgarno sequence discussed in Fig. 2 caption).

REEFFIT analysis gave excellent fits to these *add* riboswitch M^2^ data (Fig. 2n), automatically detecting the presence of P1 (14±10%), P2 (37±15%), P3 (86±18%), P4 without extension (69±10%), the extension P4B at lower population (36±6%), and P5 (94±18%) in the absence of adenine, in agreement with the apoA and apoB model from NMR (Fig. 2o-p); and recovering a holo state with P1, P2, P3, P4, and P5 dominating in 5 mM adenine (see Supporting Methods for treatment of ligand-bound structures). In ligand-free conditions, the REEFFIT analysis also gave several alternative helices in the P1/P2 region, including P1B and P2B (20±7%), consistent with the NMR-detected features in the apoB structure. The M^2^ data were critical in making these detections; RNAstructure calculations and use of wild type SHAPE data alone assigned negligible (<5%) probability to P1, P1B, P2B, and the possibility of P4B shortening to P4 (Fig. 2o-p). Coarse clustering of REEFFIT structures returned four states in which the NMR-modeled apoA was recovered as the cluster medioid of one state, ADD-A, and apoB was recovered as a medioid in another, ADD-B, albeit with an additional helix, P4B (Fig. 2p). Structures with alternative helices to P1, P2, P1B, and P2B in the 5’ region clustered into states ADD-C and ADD-D. The population fractions of these helices, as well as a set of P6 and P7 helices not detected in NMR experiments, were low; these features were appropriately flagged as uncertain from bootstrapping analysis (e.g., 21±15% for the most populated helix of this kind, P6). Overall, the M^2^-REEFFIT results successfully recovered NMR-detected helices for this adenine riboswitch sequence, including heterogeneous structure in the 5’ region, dynamics in the P4 region, and rearrangements on adenine addition.

### Assessing if artificial sequences exhibit complex RNA landscapes

After validation of M^2^-REEFFIT on diverse computational and experimental test cases, we used the method to estimate whether complex landscapes might arise in artificial RNA sequences without explicit design or selection. First, we analyzed the folding landscape of an imperfect RNA switch, ‘Tebowned’, that was designed to convert between two states upon flavin mononucleotide (FMN) binding (Figure 3) in early rounds of a riboswitch design puzzle in the EteRNA massive open laboratory.^32^ While the chemical mapping pattern of this RNA changed upon binding FMN, the measurements for the unbound state did not match the desired unbound structure, particularly near nucleotide A30 (red arrows in Fig. 3a); this region should have been paired but instead was measured to be reactive. *A priori,* we could not distinguish whether this discrepancy was due to an incorrect balance of the two target bound and unbound structures or if there were other unexpected structures involved. To elucidate the discrepancy, we acquired M^2^ data for the Tebowned RNA. We first used REEFFIT to fit the M^2^ data using only the two desired structures, but this fit did not capture several features, such as the exposure at A30 (Supporting Figure S7). In contrast, a global ensemble fit adequately captured these features (Figure 3a-b, and Fig. S7b). REEFFIT automatically clustered the modeled ensemble into three states: TBWN-A (58 ± 12%) and TBWN-B (20 ± 11%), matching the desired switch structures, and a third structure TBWN-C present at 22 ± 13% fraction (red arrows in Fig. 3c and Supporting Figures S8a-c). The unexpected state TBWN-C exhibits an apical loop around nucleotide A30, explaining the observed discrepancies in this region for the wild type RNA, and harbors a purine-rich symmetric loop that may be significantly stabilized compared to the energy assumed in current nearest-neighbor models.^33^ Other helices were discovered to be populated with up to 10% frequency in the analysis, but were deemed uncertain (signal-to-noise ratios less than 1) from bootstrapping analysis.

**Figure 3:**
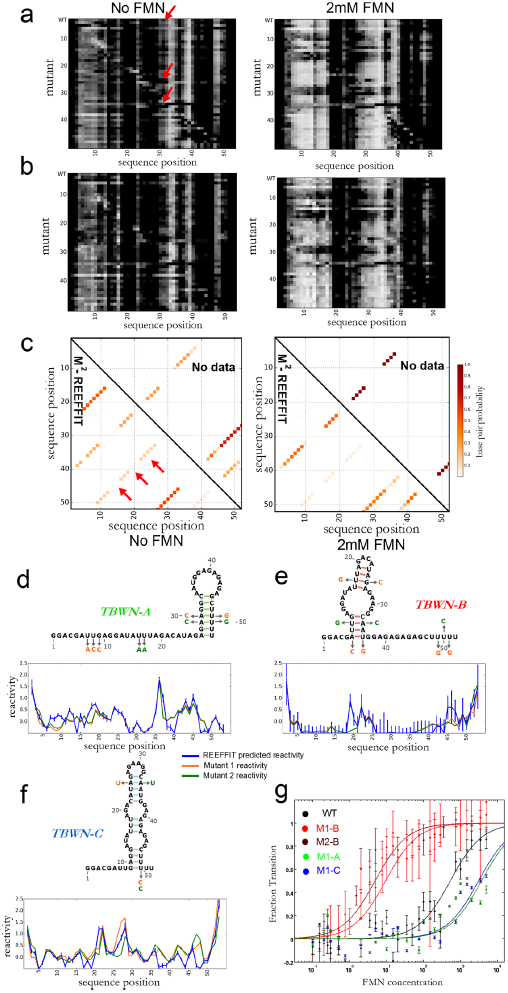
Landscape dissection of a designed FMN riboswitch with a structural discrepancy, the Tebowned RNA. (a) M^2^ data, (b) REEFFIT fit, and (c) inference of the Tebowned structural landscape, which is predicted by REEFFIT to fold into three prevalent structures given in (d) TBWN-A, (e) TBWN-B, and (f) TBWN-C (marked with red arrows in (c)). Comparison between reactivities predicted by REEFFIT and mutants designed to stabilize the TBWN-A, TBWN-B, TBWN-C structures are given below each state (arrows in the structures mark mutations). (g) Determining FMN dissociation constants for stabilizing mutants of the TBWN-A, TBWN-B, and TBWN-C structures of the Tebowned FMN switch using the LIFFT HiTRACE toolkit (showing data for residue 15, see Supporting Figure S9) are shown below the mutants.

The REEFFIT-inferred populations of TBWN-C and TBWN-C were low (<25%). Therefore, to further test the presence of at least three states, we sought to compare their modeled REEFFIT reactivity profiles to actually measured reactivities for these states. To achieve this comparison, we designed mutants that specifically stabilized helices in each of the TBWN-A, TBWN-B, and TBWN-C structures (Fig. 3d-f). For each state, the reactivities of these state-isolating mutants agreed with each other within experimental error and were approximated well by REEFFIT’s predicted profile, providing independent confirmation of the modeled ensemble, including the unanticipated third state TBWN-C. Additional evidence for the accuracy of REEFFIT’s predictions was revealed by each of the stabilizing mutants’ FMN binding affinities: the TBWN-B and TBWN-A/TBWN-C mutants enhanced and worsened ligand binding, respectively, as expected (Figure 3g, and Supporting Results and Supporting Figure S9). We emphasize that the TBWN-C state would have been difficult to propose and then validate without automated REEFFIT analysis, given its negligible predicted population in secondary structure prediction calculations without M^2^ data (Fig. 3c).

As a second test case with a previously unknown structural ensemble, we tested whether randomly generated or scrambled RNA sequences tend to fold into multiple disparate structures at equilibrium - a long-standing hypothesis fundamental to understanding RNA evolution, put forth by several *in silico* studies^34,35^ and an experimental study that could not deconvolve the structures.^36^ We carried out M^2^-REEFFIT for a randomly generated sequence, called here the M-stable RNA (Figure 4). Based on simulations, the structural ensemble of the construct was expected to consist mainly of a simple hairpin (P1 in MST-A in Figure 4d, see top triangle of the Fig. 4c) but with at least two other structures becoming more stable than MST-A upon single mutations. The experimental M^2^ measurements were complex, with different mutants giving disparate protection patterns even in segments that appeared highly reactive in the wild type RNA. As a first check on the number of states, REEFFIT fits assuming only 2 or 3 states missed many features observed in the data, including extended segments of changed chemical reactivity in several mutants (Supporting Figure S10). However, the REEFFIT global ensemble fit successfully modeled the M-stable data and suggested an ensemble with many more weakly populated helices than RNAstructure’s estimate (compare bottom and top halves of Figure 4c). For visualization, we clustered these heterogeneous component structures into three states, MST-A, MST-B, and MST-C (see Fig. 4d-f and Supporting Figures S8d-f). Analogous to the case of the Tebowned switch, we tested the REEFFIT prediction of these alternative states by designing mutations to stabilize the medioid structures of each cluster (Figure 4d-f). These mutants gave reactivities in agreement with predictions for MST-A and MST-C, supporting the inference of those structures. For MST-B, the state-stabilizing mutants gave reactivity profiles that did not exactly match each other, suggesting residual heterogeneity of structure; the profiles were nevertheless closer to the REEFFIT-predicted MST-B profile than wild type reactivities. These results corroborate the M^2^-REEFFIT model that the M-stable random RNA has a complex landscape with at least three structures, and likely significantly more heterogeneity, detectable upon unbiased nucleotide mutation.

**Figure 4:**
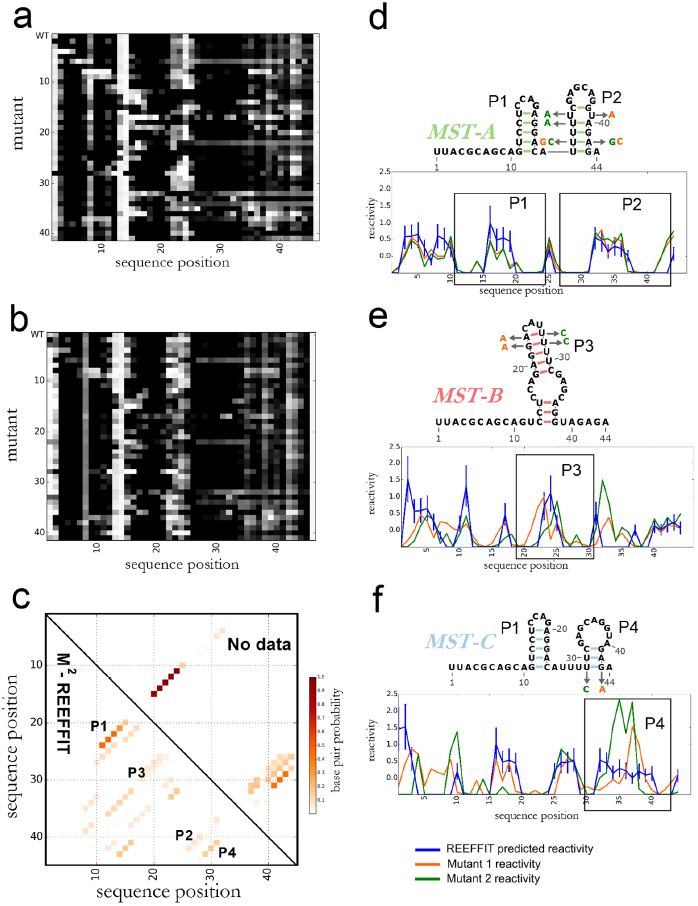
Landscape dissection of a random RNA sequence, the M-stable RNA. (a) M^2^ data, (b) REEFFIT fit, and (c) inference of the M-stable structural landscape, showing one helix, P1, present at almost 50% as well as numerous other helices (P2, P3, and P4) present at non-negligible fractions. (d-f) State medioids for (d) MST-A, (e) MST-B, and (f) MST-C, and comparison between reactivities predicted by REEFFIT and mutants designed to stabilize each state medioids.

## DISCUSSION

The structure landscapes of natural and newly designed RNA molecules underlie their biological behaviors, but these landscapes’ complexities are largely uncharacterized.^11^ Current experimental techniques used to probe these ensembles at nucleotide resolution require significant infrastructure investment and expert intuition. We have presented M^2^-REEFFIT, an unbiased strategy based on readily acquired chemical mapping measurements that detects dominant and alternative states of RNA structure landscapes in the ensemble perturbations induced by single-nucleotide mutations, with conservative estimates based on bootstrapping.

Due to the challenges inherent to estimating a full ensemble of secondary structures rather than a single best-fit model, we have invested significant effort into benchmarking M^2^-REEFFIT and its uncertainty estimates. We confirmed the accuracy of REEFFIT and its uncertainty estimates in M^2^ simulation data with available ‘ground truth’ landscapes and in experimental data of RNAs whose landscapes were previously studied by chemical mapping or NMR experiments. We then applied REEFFIT to investigate if artificial sequences could present complex landscapes of alternative structures without specific design in two model systems whose ensemble behaviors were refractory to prior tools. We first traced a structural discrepancy in an artificial flavin-mononucleotide-binding switch to a significant population of an unexpected state. We then tested *in silico* predictions for the complex structural landscape of random sequences by obtaining the first experimentally derived landscape model of a randomly generated sequence, the M-stable RNA. For all test cases, we tested REEFFIT’s predictions through independent experiments, including comparison to independent experimental methods and effects of stabilizing mutations and ligand binding.

Currently, uncertainties in the reactivities and energies of RNA motifs lead to RMSD errors in M^2^-REEFFIT state population fractions on the order of 10%, rendering the detection of states with lower population fractions difficult; these uncertainties may become poorer for longer RNA domains. Nevertheless, we expect that rapidly growing databases of rigorously standardized reactivity data^27,37^ and of energetic parameters^38^ for diverse RNA motifs will reduce these uncertainties. Through the presented and related chemical mapping technologies, we therefore expect to have more routine visualization of the rich structural landscapes that appear to be pervasive in both functional RNAs and the generic sequences from which they evolve.

## METHODS

### M^2^ measurements

The MedLoop RNA mutate-and-map data were obtained as part of the experimental pipeline of the EteRNA massively parallel open laboratory^32^ and were analyzed with the MAP-seeker software.^39^ The Bistable RNA and its complementary single-nucleotide mutants were constructed using PCR assembly, in vitro transcription, and probed with 1M7 as described previously.^40^ Briefly, an assembly consisting of at most 60-nucleotide primers was designed to synthesize an in vitro transcription sample by PCR. DNA was purified with AMPure XP beads (Agencourt, Beckman Coulter) and in vitro transcribed for 3 hours. The resulting RNA was purified with AMPure XP beads, heated for 3 minutes at 90 °C, cooled at room temperature for 15 minutes, and folded in 150 mM NaCl at room temperature for 1 hour. Because we wanted to probe the ensemble of the RNA with minimal interference from the 3’ unpaired sequence that we use as the primer binding site, we folded the RNA in the presence of the fluorescent primer attached to the oligo(dT) beads (Ambion) that we regularly use for purification. Folding in this condition sequesters any additional single stranded regions that may interfere with our sequence of interest.. The RNA was then subjected to 1M7 mapping (5 mM final concentration), purified with the oligo(dT) beads, and reverse transcribed for 30 minutes at 42°C. Unmodified RNA controls were also included in the experiment. RNA was then degraded using alkaline hydrolysis and cDNA was purified, eluted in Hi-Di Formamide spiked with a fragment analysis ladder (ROX 350 standard, Applied Biosystems), and electrophoresed in an ABI 3150 capillary electrophoresis sequencer. The add adenine riboswitch M^2^ data was obtained similarly but folded under different conditions, in the NMR buffer of the previous study (50 mM KCl and 25 mM K_3_PO_4_, pH 6.5) with or without 5 mM adenine for bound and unbound conditions - these conditions also included 10 mM MgC_2_^31^ For NMR folding conditions, we adjusted the 1M7 incubation time to 15 minutes instead of 3 minutes to account for the low pH. The Tebowned RNA M^2^ data was obtained using the same protocol, but with a different folding condition, 50 mM Na-HEPES, pH 8.0 and 10 mM MgCl_2_, in the absence and presence of 2 mM FMN. For the Tebowned FMN titrations, dimethyl sulfate (DMS) was used in lieu of 1M7 since it yields a readily seen signal change across FMN concentrations.^32^ The M^2^ data of the M-stable RNA was obtained in a similar manner, folding the RNA by heating and cooling in 50 mM Na-HEPES, pH 8.0 and letting it fold in 10 mM MgCh. (see also Supporting Table S9 for a list of folding conditions).

Electrophoretic traces were aligned, baseline subtracted, and normalized with the HiTRACE MATLAB toolkit.^41^ 1M7 modification traces were quantified, background subtracted, and corrected for attenuation using 10X dilutions, the unmodified controls, and the pentaloop hairpins added at the ends of the constructs as reference.^42^ For the Tebowned RNA, no pentaloop hairpins were added and we relied instead on the HiTRACE background subtraction routine overmod _and_background_correct_logL using unmodified controls. The lifft function from the LIFFT package^43^ in HiTRACE was used to calculate the FMN binding dissociation constants for REEFFIT comparisons. Mutants with low signal were flagged as low quality and were not taken into account for the analysis.

### The RNA Ensemble Extraction From Footprinting Insights Technique (REEFFIT)

For a given mutate-and-map data set, REEFFIT infers the expected reactivity profile of each structure, the combination of structure population fractions (also denoted here as structure weights), and sequence-position-wise noise levels that best fit the data (see Supporting Figure S1) using prior information on known chemical reactivity distributions, a prior based on an approximate secondary structure energetic model, and a well-defined likelihood function. We present the basic idea behind the model below. Detailed description of the priors and parameters used for each variable, their incorporation into the model, and inferential steps used to fit the model to the data are given in the Supporting Methods of the Supporting Information.

In a broad sense, for *m* chemical mapping measurements of *n*-nucleotide sequences, REEFFIT models the data, denoted as *D*^*obs*^ ∊ ℜ^m×*n*^, with a set of *r* secondary structures. The data are modeled as linear combinations of the structures’ reactivity profiles, denoted as a matrix *D*∊ ℜ ^r×*n*^, with a weight matrix *W* ∊ ℜ^m×*r*^ plus Gaussian noise. Then, for each measurement *j* we have:

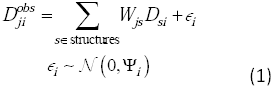

The rows of the weight matrix *W* correspond to the population fractions of the structures in each measurement. Therefore, the corresponding weights for each measurement define a probability distribution and are non-negative and add up to one:

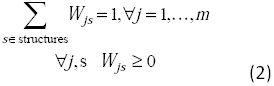

The model including these variables and parameters was fitted by REEFFIT using a maximum a posteriori (MAP) approximation.

## ACKNOWLEDGMENTS

We thank W. Kladwang, J. Lee, A. Treuille, and B. Seo for implementing mutate-and-map measurements on the EteRNA platform; E. Ashley for computational resources; F. C. Chou for careful comment and critique; N. Bisaria, B. Alford, T. Mann, members of the Das lab, and the EteRNA player community for comments and useful discussion. This work was funded the Stanford Bio-X Interdisciplinary Initiative Program, Stanford Media-X, a CONACyT graduate student fellowship (to PC), a Burroughs-Wellcome Foundation Career Award (to RD), the Keck Medical Research Foundation, and National Institutes of Health Grant R01 R01GM100953.

